# Evidence for widespread positive and negative selection in coding and conserved noncoding regions of *Capsella grandiflora*

**DOI:** 10.1101/002428

**Authors:** Robert J. Williamson, Emily B. Josephs, Adrian E. Platts, Khaled M. Hazzouri, Annabelle Haudry, Mathieu Blanchette, Stephen I. Wright

**Affiliations:** Department of Ecology and Evolutionary Biology, University of Toronto 25 Willcocks St, Toronto ON M5S 3B2; Centre for Bioinformatics, McGill University, Montreal, Quebec, Canada; School for Computer Science, McGill University, Montreal, Quebec, Canada; Center for Genomics and Systems Biology, New York University Abu Dhabi, Abu Dhabi, United Arab Emirates; Université Lyon 1, Centre National de la Recherche Scientifique (CNRS), Unité Mixte de Recherche (UMR) 5558, Laboratoire de Biométrie et Biologie Evolutive, Villeurbanne, France.; Centre for the Analysis of Genome Evolution and Function, University of Toronto, Toronto, Ontario, Canada

## Abstract

The extent that both positive and negative selection vary across different portions of plant genomes remains poorly understood. Here, we sequence whole genomes of 13 *Capsella grandiflora* individuals and quantify the amount of selection across the genome. Using an estimate of the distribution of fitness effects, we show that selection is strong in coding regions, but weak in most noncoding regions, with the exception of 5’ and 3’ untranslated regions (UTRs). However, estimates of selection in noncoding regions conserved across the Brassicaceae family show strong signals of selection. Additionally, we see reductions in neutral diversity around functional substitutions in both coding and conserved noncoding regions, indicating recent selective sweeps at these sites. Finally, using expression data from leaf tissue we show that genes that are more highly expressed experience stronger negative selection but comparable levels of positive selection to lowly expressed genes. Overall, we observe widespread positive and negative selection in coding and regulatory regions, but our results also suggest that both positive and negative selection in plant noncoding sequence are considerably rarer than in animal genomes.

**Author Summary:** Selection affects patterns of genomic variation, but it is unclear how much the genomic effects of selection vary across plant genomes, particularly in noncoding regions. To determine the strength and extent of selective signatures across the genome, we sequenced and analyzed genomes from 13 *Capsella grandiflora* individuals. *C. grandiflora* is well-suited for this task because it has experienced a large, stable effective population size, so we expect that selection signatures will not be overly distorted by demographic effects. Our analysis shows that positive and negative selection acting on new mutations have broadly shaped patterns of genomic diversity in coding regions but not in most noncoding regions. However, when we focus only on noncoding regions that show evidence of constraint across species, we see evidence for strong positive and negative selection. In addition, we find that genes with high expression experience stronger negative selection than genes with low expression, but the extent of positive selection appears to be equivalent across expression categories.

## Introduction

Determining the amount of positive and negative selection and how it varies across the genome has wide-ranging implications for understanding genome function and the maintenance of genetic variation [1]. Current evidence suggests that both positive and negative selection are common in coding and some noncoding sequences in several model systems [2-8]. However our understanding of genome-wide selection in plants remains relatively limited [6], particularly in noncoding regions.

The extent to which both positive and negative selection operate in noncoding regions of the genome compared with coding regions remains unclear [2,5-8]. For example, it has been suggested that the majority of adaptive evolution may occur in noncoding regulatory regions, where new mutations may have fewer deleterious pleiotropic effects [9,10 but see 11]. Halligan and colleagues [8] showed that there have been many more adaptive substitutions in noncoding DNA than in coding regions in house mice, although adaptive substitutions in coding regions may experience stronger positive selection. Moreover, studies in *Drosophila* and vertebrates have found that, although noncoding regions as a whole are generally less conserved than coding regions, there is more functional noncoding sequence than constrained coding sequence by a considerable margin [2,12].

Comparing these results to noncoding selection across plant genomes is of particular interest because it has been hypothesized that in plants, regulatory evolution may occur more often through gene duplication than *cis-*regulatory change [13], possibly leading to lower levels of functional constraint and positive selection in plant noncoding DNA. Consistent with this prediction, Haudry and colleagues [14] recently compared the genomes of nine Brassicaceae species, and showed that approximately one quarter of the conserved sites in the *A. thaliana* genome were in noncoding regions, a much smaller fraction than found to date in studies of vertebrates and *Drosophila*. However, the strength of selection on these noncoding sites, the extent of species-specific selection on noncoding regions, and the extent of positive selection in noncoding regions compared with coding regions have not been quantified.

While the strength of selection is expected to vary between coding and noncoding sequence, it also varies between genes. Gene expression level is one of the major determinants of rates of nonsynonymous evolution in coding regions in many species [15-17], including plants [18-21]. Variation in the strength of selection on genes could reflect differences in the relative importance of gene products for organism fitness, or it may simply relate to inherent properties of expression [22]. For example, deleterious mutations causing misfolding or mis-interaction have more opportunity to interfere with cellular function when they occur in high expression genes [23-25]. Regardless of the underlying selective mechanisms, the negative correlation between expression level and nonsynonymous divergence could reflect relaxed purifying selection in lowly expressed genes, increased positive selection on lowly expressed genes, or both.

Here, we use population genomics to quantify the strength of both positive and negative selection inside and outside of coding regions and within highly and lowly expressed genes in a species-wide sample of 13 outbred *Capsella grandiflora* individuals. *C. grandiflora* is an obligately outcrossing member of the Brassicaceae family with a large effective population size (*N_e_* ∼ 600,000) and relatively low population structure [26,27]. We estimate the strength of negative selection by fitting polymorphism data to a model of the distribution of negative fitness effects of mutations. We then quantify the contribution of positive selection to divergence in *C. grandiflora* using two complimentary approaches: an extension of the McDonald-Kreitman test [28] and an analysis of neutral variation linked to lineage-specific fixed substitutions [29]. Our results demonstrate that both positive and negative selection are pervasive in coding regions, 5’ and 3’ untranslated regions (UTRs), and constrained noncoding regions of the *C. grandiflora* genome, but also that a large proportion of noncoding DNA may evolve neutrally. In addition, we find stronger negative selection in high expression genes compared to low expression genes, suggesting that differences in negative selection drive differences in rates of molecular evolution.

## Results

### Genome wide patterns of polymorphism

We sequenced 13 outbred *C. grandiflora* individuals (26 sampled haploid chromosomes; ∼140 Mb genome assembly) sampled from across the species’ range in northern Greece using Illumina GAII sequencing (Supplementary Table S1). The resulting 108 bp reads were mapped to the *C. rubella* reference genome [30] using the Stampy aligner resulting in a median coverage of 34 reads per sample per site. Genotypes were called using the Genome Analysis Toolkit’s Unified Genotyper [31]. After filtering for quality and depth (see Methods), we were left with 27,034,975 sites, 1,538,085 of which were single nucleotide polymorphisms (SNPs) (Table S2). Sites from across the genome were identified as 0-fold degenerate, 4-fold degenerate, intronic, 5’ UTR, 3’ UTR, or intergenic, based on the annotation of the *C. rubella* reference genome [30]. To avoid comparing sites that would not have equivalent mutation profiles, we excluded sites in coding regions that were neither 4-fold nor 0-fold degenerate. After filtering, our analysis includes 30-40% of coding and noncoding sites, except in intergenic regions where only approximately 10% of sites are retained due to the higher repeat content in these regions and the removal of highly repetitive pericentromeric DNA (Supplementary Figure S1).

Consistent with previous estimates made using a much smaller set of loci (257 Sanger-sequenced loci) and a different range-wide sample [32], average nucleotide diversity at 4-fold degenerate sites was 0.022 and there was evidence for an excess of rare variants genome-wide at 4-fold degenerate sites compared with the standard neutral model (Tajima’s *D* = -0.512). Introns (Watterson’s *θ_w_* = 0.020) and intergenic regions (*θ_w_* = 0.019) showed only slightly lower levels of nucleotide diversity than 4-fold degenerate sites, suggesting that the large majority of sites in these regions are effectively neutral, or subject to comparable levels of purifying selection as 4-fold degenerate sites. 5’ and 3’ UTRs showed a much stronger diversity reduction (*θ_w_* = 0.015 and 0.014 respectively), while 0-fold degenerate nonsynonymous sites showed the strongest reduction (*θ_w_* = 5).

Neutral diversity at 4-fold degenerate sites near centromeric regions was elevated on most chromosomes, similar to observations made in *Arabidopsis thaliana* [33], *Arabidopsis lyrata* [34,35] and *Medicago truncatula* [36], (Supplementary Figure S2). As with these other species, this effect is not obviously caused by higher mutation rates in these regions, since divergence between *Capsella* and *Neslia* is not clearly elevated in these regions (Supplementary Figure S2). Although elevated error rates in repetitive regions may contribute to high diversity, our observation of high diversity in these regions is still apparent after extensive filtering (see Methods). This increase in neutral diversity in pericentromeric regions may reflect a weakening of background selection in regions of low gene density, as recently shown in models of background selection applied to *Arabidopsis* [37]. Furthermore, diversity generally declines towards the ends of the chromosomes, potentially reflecting the stronger effects of background selection and/or selective sweeps in regions of relatively low recombination but high gene density, where the effects of linked selection are expected to be strongest. Consistent with these interpretations, we see an increase in diversity in regions of low coding density (Supplementary Figure S3).

We also examined individual heterozygosity in sliding windows along each chromosome. A number of individuals showed large stretches of homozygosity indicative of biparental inbreeding (Supplementary Figure S4 and Supplementary Figure S5). Consistent with these regions reflecting local biparental inbreeding, no such regions are found in our sample that is derived from a between-population cross, AXE. These regions of identity-by-descent (IBD) in our data highlight that, despite being self-incompatible and obligately outcrossing, local biparental inbreeding can still generate excess homozygosity in stretches across the genome. To avoid biased estimation of species-wide allele frequencies in these regions, we subsampled the data to treat all IBD regions as haploid rather than diploid sequence for the purposes of allele frequency estimation, although treating these regions as diploid does not qualitatively change our conclusions (Supplementary Figure S6).

### Genome-wide measures of purifying selection

Using the methods of Eyre-Walker & Keightley [1] we compared the allele frequency spectrum (AFS) and divergence for putatively neutral sites (4-fold degenerate) to those from every other category of sites in order to quantify the amount of negative and positive selection on each category (Figure 1A). Consistent with the patterns of diversity described above, negative selection is generally much stronger in coding regions than noncoding regions (Figure 1A). This pattern is most clearly seen in 0-fold degenerate sites, the only site category with a sizable fraction of sites in the strongest category of negative selection (41%). Of the noncoding categories, UTRs show much stronger negative selection than other regions. In *C. grandiflora* ∼55% of both 5′ and 3′ UTRs are under moderate levels of purifying selection (*N_e_s* > 1), but a considerably larger fraction of UTR sites are effectively neutral (45%) than 0-fold degenerate sites (14%). Additionally while the UTRs and CNSs (see below) show a signal of strong purifying selection (*N_e_s*> 10) the amounts of strong selection on these sites is lower than in 0-fold degenerate sites.

**Figure 1.**
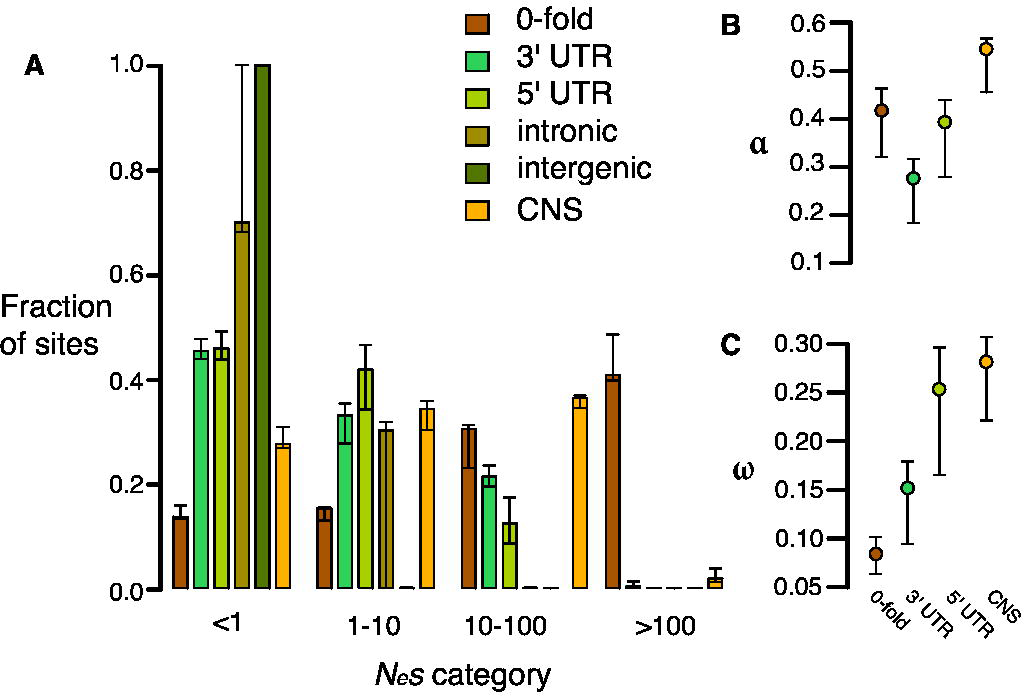
Estimates of negative and positive selection on coding and noncoding sites in *C. grandiflora*. A) The proportion of sites found in each bin of purifying selection strength, separated by site type, B) The proportion of divergent sites fixed by positive selection, and C) the rate of adaptive substitution relative to neutral divergence. Error bars represent 95% bootstrap confidence intervals.

Genome-wide, we estimate that the proportion of intergenic sites that are nearly neutral approaches 100% and that approximately 70% of intronic sites are effectively neutral. Furthermore, bootstrapping results suggest that there is not significant support for less than 100% of intronic sites being effectively neutral. The large confidence intervals around estimates of selection on intronic sites may be due to strong selection at splice site junctions [14] coupled with typically weak to no selection outside of splice junctions. To test for selection near splice junctions, we quantified selection acting on the first and last 30 bp of each intron separately from sites in the middle of introns. While 100% of sites in the middle of introns are estimated to be effectively neutral, only 68% of sites in junctions are, suggesting that our wide confidence intervals around intronic sites can be partially explained by variance caused by sampling sites in these different regions between bootstraps. These generally low estimates of *N_e_s* in (non-junction) intronic and intergenic sites imply a general lack of purifying selection in most noncoding regions, a lack of sensitivity to detect small proportions of selected sites, and/or nearly equivalent purifying selection to synonymous sites.

Although our analysis suggests very low levels of purifying selection in noncoding regions other than UTRs and splice junctions, these global analyses may miss signatures of purifying selection on a small proportion of noncoding sites. One candidate set of sites that may have different signatures of selection are conserved noncoding sequences (CNSs); these are regions that show evidence of cross-species conservation, and are therefore prime candidates for functional noncoding sequences subject to selection. We identified CNSs across nine Brassicaceae genomes, following the implementation in Haudry et al. [14]. For this study, we used the *Capsella* genome as a reference for alignment, but excluded *Capsella* when identifying CNSs in order to avoid circularity when quantifying selection from diversity [8]. This method allows our analysis of selection on noncoding sites using polymorphism to be more independent of the comparative analysis. When we look at only these conserved regions in our *C. grandiflora* sample we see a small proportion of effectively neutral sites (28%) compared to the noncoding regions as whole, suggesting that the majority of CNS sequences are subject to purifying selection (Figure 1). However, estimates suggest that CNSs are generally under weaker purifying selection than nonsynonymous (0-fold) sites and experience primarily weak and intermediate purifying selection (Figure 1).

Although CNSs as a whole retain a considerable proportion of effectively neutral sites, it is of interest to examine whether particular classes of CNS show stronger selection. To examine differences between categories we quantified selection on the different types of CNSs separately (Supplementary Figure S7). In most categories, about 25% of sites are nearly neutral, a slightly stronger signal of purifying selection than when we pool all CNSs. Intronic CNSs have a larger proportion of effectively neutral sites than other categories, in agreement with the general neutrality of intronic sites (Figure 1). In contrast, small noncoding RNAs (sncCNSs) have a stronger signal of selection than the other CNS categories. However, the number of sites used to make the AFS for each of these categories varies substantially (Supplementary Table S2), and our sample of sncCNSs has very little polymorphism (155 segregating sites). Nevertheless, despite the wide confidence intervals, sncCNSs still show a significantly (*p < 0.001*) smaller fraction of sites that are nearly neutral (*N_e_s* < 1) than the pooled CNSs, which could be due to strong selection for sequence specificity to obtain the proper secondary structure important for RNA activity [38]. This effect is consistent with sncCNSs showing a higher degree of conservation across the Brassicaceae [14] and having traceable orthologs in other plants.

### Genome-wide estimates of positive selection

We used the approach of Eyre-Walker and Keightley [28] to estimate the proportion of fixations driven by positive selection (α) and the rate of positive selection (ω) while taking into account the effect of slightly deleterious mutations, which can bias estimates of positive selection downwards. To do this, we estimated divergence using whole genome alignments of *Capsella*, *Arabidopsis thaliana*, and *Neslia paniculata* (estimate of 4-fold synonymous divergence *K_s_* between *C. rubella* and *N. paniculata* is *K_s_*=0.14). Because the large majority of noncoding sites are estimated to be effectively neutral, and because of alignment concerns between species in unconstrained noncoding regions, we focus our estimates of positive selection in noncoding DNA on CNS sites and UTRs. We found that 0-fold degenerate sites show a very high proportion of divergence driven by positive selection (Figure 1B; α = 0.417) and estimates of the rate of adaptive substitution relative to synonymous substitution (Figure 1C; ω = 0.08). Similarly, UTRs and CNS sites show evidence for positive selection (Figures 1B and 1C). These results generally suggest widespread positive selection on both nonsynonymous and functional noncoding genomic regions.

If many of the amino acid changes between *C. grandiflora* and its nearest relatives are due to recent, strong positive selection from new mutations, we expect to see the signature of selective sweeps: reduced neutral diversity surrounding amino acid fixations [39,40]. We tested for this signature by measuring the proportion of 4-fold degenerate sites in each window that were polymorphic (referred to subsequently as ‘4-fold diversity’) in non-overlapping 1 kb windows surrounding fixed replacement (n = 60,378), and silent (n = 83,812) substitutions in *C. grandiflora*. 4-fold diversity surrounding fixed replacement substitutions was lower than 4-fold diversity surrounding fixed silent substitutions in the 4kb window surrounding substitutions (Figure 2A). This result was robust to various window sizes from 500kb to 2kb (Supplementary Figure S8) and a one-tailed test for reduced 4-fold diversity around replacement sites was significant (*p < 0.01* for 2 kb on either side of the substitution).

**Figure 2.**
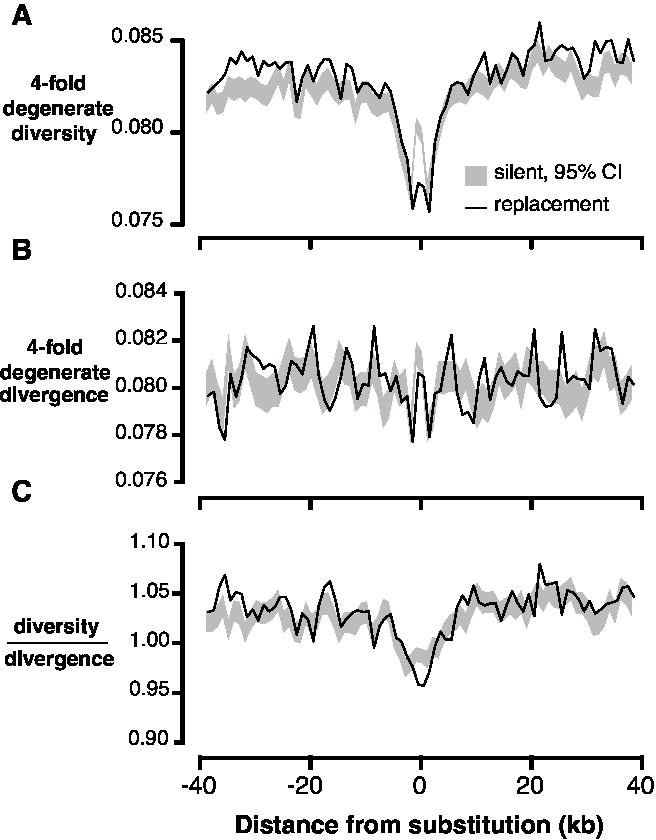
Linked neutral diversity and divergence as a function of distance from fixed substitutions across the *C. grandiflora* genome. A) 4-fold degenerate diversity, B) 4-fold degenerate divergence, and C) 4-fold degenerate diversity/divergence. In all figures, black lines represent measures surrounding fixed replacement substitutions and gray shading represents 95% confidence intervals, from bootstrapping, surrounding silent substitutions.

Patterns of diversity may be distorted by elevated mutation rates surrounding substitutions [39], which would increase diversity and divergence in *C. grandiflora*. Consistent with this prediction, divergence at 4-fold degenerate sites (‘4-fold divergence’) is elevated around synonymous and replacement substitutions (Figure 2B). To control for elevated mutation rate, we divided diversity by divergence at 4-fold degenerate sites (subsequently referred to as ‘4-fold diversity/divergence’). We observed a reduction in 4-fold diversity/divergence around replacement substitutions compared to silent substitutions, demonstrating that the signature of recurrent sweeps is not an artifact caused by variation in mutation rate (Figure 2C, *p < 0.01* for 1 kb on either side of the substitution).

An analogous test for selective sweeps around fixations in noncoding regions is challenging because the test depends on accurately identifying interspersed functional and neutral sites, a difficult task in noncoding regions [8]. Instead, we compared 4-fold diversity and divergence around fixed substitutions in CNS regions (n = 12,578) with 4-fold diversity and divergence around fixed substitutions in non-conserved intergenic, intronic, and UTR regions (n = 117,178). Interestingly, there is a reduction in both 4-fold diversity and divergence surrounding fixed substitutions in CNSs compared to non-conserved noncoding regions (Supplementary Figure S9). It is not clear why 4-fold divergence decreases around CNS substitutions; it is possible that in genomic scans for conserved regions, large-scale constraint might span both coding and noncoding sequence, causing non-independence and reducing divergence at 4-fold degenerate sites near CNSs. However, there is still a reduction in 4-fold diversity/divergence around fixed substitutions in CNSs compared to those in non-conserved intergenic regions, consistent with the action of recurrent selective sweeps (Figure 3A).

**Figure 3.**
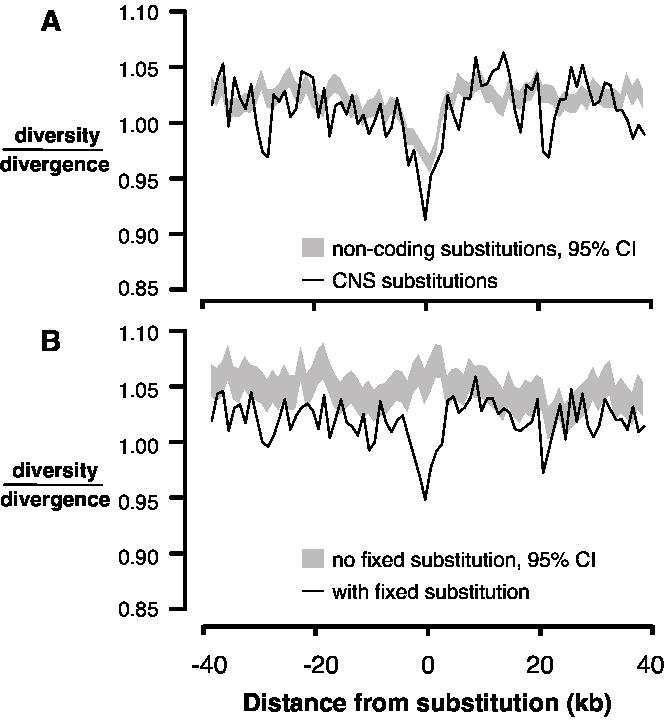
Linked neutral diversity/divergence surrounding conserved noncoding sequences (CNSs). A) 4-fold degenerate diversity/divergence as a function of distance from fixed substitutions in CNSs (black lines) and fixed substitutions in non-conserved intergenic sequence (gray shading, 95% confidence interval). B) 4-fold degenerate diversity/divergence as a function of distance from CNSs containing fixed substitutions (black line) and CNSs without any fixed substitutions (gray shading, 95% confidence interval).

The observed reduction in diversity/divergence around CNS substitutions could also reflect the action of background purifying selection; sites closer to CNSs may experience a reduction of neutral diversity due to greater purifying selection on mutations in CNSs. This effect is not a problem for comparisons between replacement and silent substitutions because they are interspersed within the same exons, so diversity and divergence around these sites experience the same background selection. To ensure that the reduction in diversity/divergence surrounding CNS substitutions compared to non-conserved noncoding substitutions is not due to differences in background selection between CNS and intergenic sites, we compared neutral diversity and divergence surrounding CNSs that contain at least one fixed substitution to neutral diversity and divergence around those that do not. There is a reduction in neutral diversity/divergence surrounding CNSs containing a fixed substitution (n = 12,884) compared to CNSs without fixed substitutions (n = 41,212), suggesting that this signature of recurrent sweeps is not driven by background selection specific to CNSs (Figure 3B).

### Effects of expression and selection

We measured expression levels of all expressed genes using RNA extracted from leaf tissue of 10 of the 13 *C. grandiflora* individuals. Genes were sorted by mean expression level and split into four equally sized groups, which will be referred to as “high”, “mid-high”, “mid-low”, and “low” expression genes. We calculated polymorphism within *C. grandiflora* and lineage-specific divergence from *N. paniculata* and *A. thaliana* for sites within these genes. As expected from previous studies, *dN*/*dS* is considerably lower in high expression genes (0.15) than low expression genes (0.22). In addition, *dN/dS* is negatively correlated with expression level across all genes (correlation coefficient = −0.051, *p < 0.001*).

To test whether the strength of negative selection differs between expression categories we compared the allele frequency spectra of sites in different expression categories. Replacement polymorphisms in high expression genes show a stronger skew towards rare alleles than those in low expression genes (Supplementary Figure S10). In addition, a larger proportion of replacement sites are invariant in high expression genes (98.9%), than in low expression genes (97.8%), consistent with stronger negative selection. The distribution of fitness effects, estimated using a randomization test [28], show that high expression genes have a much smaller proportion of effectively neutral sites (6.8%) than low expression genes (16%, *p < 0.001*) (Figure 4A).

**Figure 4.**
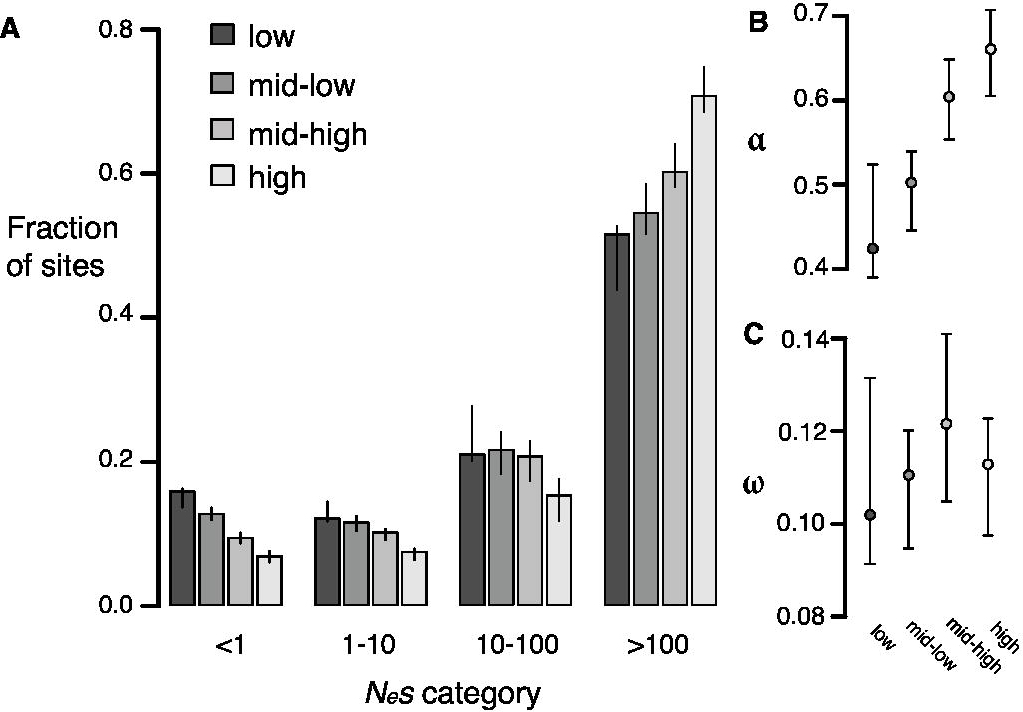
Estimates of negative and positive selection on 0-fold degenerate sites in genes of varying expression level. A) The proportion of sites found in each bin of purifying selection strength, separated by site type. B) The proportion of divergent sites fixed by positive selection and C) The rate of adaptive substitution relative to neutral divergence. Error bars represent 95% bootstrap confidence intervals.

Increased divergence in low expression genes relative to high expression genes could also be caused by increased positive selection in low expressed genes compared to highly expressed genes. To test this possibility, we calculated α and ω as described above. High expression genes have a significantly higher value of α (0.66) than low expression genes (0.42, *p < 0.01*) but the ω value for both classes is similar (high: 0.11, low: 0.10, *p = 0.38*), suggesting that the rate of positive selection does not differ between high and low expression genes (Figure 4B, 4C). The difference in α between the two categories likely reflects the reduction in the number of weakly deleterious and effectively neutral mutations that are able to fix due to stronger purifying selection in high expression genes compared to low expression genes, causing a higher proportion of those amino acids that do reach fixation to be positively selected.

## Discussion

In this population genomic survey of *C. grandiflora*, we demonstrated that positive and negative selection contribute to DNA sequence variation in protein-coding regions, UTRs, and CNSs. We also showed that differences in divergence between high and low expression genes are due to increased negative selection on high expression genes, not increased positive selection on low expression genes. In addition, we found a clear signature of recurrent selective sweeps contributing to divergence in coding regions as well as CNSs. Overall, our evidence for widespread positive and negative selection in *C. grandiflora* is in line with expectations, given its outcrossing mating system, large *N_e_*, limited population structure, and lack of a recent whole genome duplication [6].

In contrast, selection appears to be very rare in intergenic and (non-junction) intronic regions that are not conserved across *Brassicaceae* species. In particular, we cannot detect significant evidence of purifying selection on intergenic or intronic regions as a whole, suggesting that selected sites within these regions must be rare or absent. However, when we only examine CNSs, we do see evidence of selection, indicating that at least 5% of sites in intergenic regions are selected, but the DFE approach is not sensitive enough to detect selection on such a small proportion of the genome. This result implies that this approach is likely to also be missing lineage-specific selection when it comprises a relatively small fraction of sites, and it highlights the importance of integrating additional evidence of function (comparative and experimental) for improved quantification of selection.

The general neutrality of noncoding regions, based on population genomic analysis, is consistent with the conclusions of Haudry and colleagues [14], who used comparative genomics approaches to estimate that only 5% of noncoding bases are under selection in the *Arabidopsis* genome. This result contrasts with *Drosophila* and humans, where a relatively large fraction of selected sites are found in noncoding regions [6]. For example, in *Drosophila*, only 30%-70% of intronic and intergenic regions are nearly neutral (Andolfatto 2005, Sella 2009, Eyre-Walker & Keightley 2009). Similarly, Halligan et. al. [8] recently used information from the DFE to infer the number of adaptive substitutions in mice both in coding and noncoding regions. They show that the majority (approximately 80%) of the adaptive substitutions in the mouse genome are in noncoding regions and suggest that they may have regulatory function. In contrast, our data show that *C. grandiflora* has similar numbers of adaptive substitutions in 0-fold sites (50.6 kb) and noncoding sites (21.6 kb, 3’ UTR excluding CNSs; 10.2 kb, 5’ UTR excluding CNSs; 32.7 kb, CNS; 64.4 kb total). Additionally, the width of diversity reductions surrounding replacement substitutions and substitutions in CNS regions appear comparable, suggesting that there is little evidence for a difference in the strength of positive selection on substitutions in coding regions compared to conserved noncoding regions. Our results are consistent with previous suggestions that, unlike in animals, plant genomes may contain fewer noncoding regulatory sequences subject to positive and negative selection, possibly because gene expression can be modified through frequent gene duplication and functional divergence rather than through the evolution of novel regulatory elements [13].

Unlike other classes of noncoding sequence, UTRs show relatively high levels of purifying selection, likely reflective of their function in post-transcriptional regulation [41]. UTRs are also under stronger negative selection than other noncoding regions in *Drosophila* [2], and this result is also in line with the previous study using comparative genomics in the Brassicaceae [14]. Interestingly, we infer that a large fraction of selected sites in UTRs may be outside of CNS regions identified in between-species comparisons. In particular, using estimates of the proportion of sites under selection, we estimate that 88% of 3’ UTR and 77% of 5’ UTR selected sites are outside of conserved regions. This result suggests that there may be many species-specific (i.e., non-CNS) functional regions in UTRs and they may therefore play an important role in recent or local adaptation.

One important consideration is the extent to which our analyses are truly reflective of genome-wide patterns of selection. Despite whole genome sequencing, our analyses are restricted to approximately 20% of the genome, and only 10% of intergenic sites, largely due to the fact that a large fraction of the genome is pericentromeric, repetitive and/or surrounds insertion/deletion events. It is important to recognize that our estimates of selection apply strictly to this ‘accessible’ genome and that the extent of purifying and positive selection in the repetitive regions remains difficult to assess. Nevertheless, we would expect that our conclusions about low levels of purifying and positive selection across most noncoding regions is likely conservative, with respect to these filters, because a large proportion of repetitive DNA is likely to be neutral. On the other hand, rates of positive selection may be elevated in coding regions of duplicate genes filtered out of our analysis [42], suggesting that our estimates of positive selection on protein-coding regions may also be a lower bound.

A second concern is the extent to which synonymous sites are neutrally evolving. Although analysis of codon usage bias from population genetic data does suggest the action of some purifying selection on synonymous sites in this species [43], the strength of selection inferred is close to effective neutrality. Furthermore, synonymous site selection is expected to be stronger in more high expression genes [23,44], causing us to underestimate, rather than overestimate, the difference in the strength of purifying selection compared with low expression genes. Thus, while selection on synonymous sites may bias our estimates of selection slightly downward, our general conclusions are likely to be robust to violations of neutrality. Nevertheless, more investigation of the action of selection on synonymous sites is important, particularly given growing evidence for synonymous site selection that may reflect gene regulation, in addition to codon usage [45,46].

At synonymous sites, we see an excess of rare variants, as indicated by a negative Tajima’s D. The excess of rare variants is unlikely to be explained by a high Illumina error rate, as our observed value of −0.51 is nearly identical to a previous estimate (−0.52) from Sanger-sequenced loci and a comparable geographic sampling [27]. This previous study found that, while population subdivision was low compared to other herbaceous species studied, there were still three major geographic clusters (average between-population F_st_ of 0.11). If we restrict our dataset to one of the three geographic regions based on these previous results, Tajima’s D approaches zero (−0.16 at 4-fold degenerate sites), suggesting that the excess of rare variants at synonymous sites may be largely due to population structure.

### Measuring positive selection

In this study, we took advantage of the two detectable signatures expected to remain after recurrent classic selective sweeps from new mutations: 1) an excess of replacement substitutions relative to expectations based on polymorphism, and 2) reduced neutral diversity near fixed differences. Our findings strongly suggest that positive selection has been common in coding regions, UTRs and conserved noncoding regions in *C. grandiflora* and that classic selective sweeps contribute significantly to divergence in these regions. To our knowledge, this is the first time that the signature of recurrent selective sweeps has been observed in a non-*Drosophila* species, despite being tested in other species [8,47]. Our ability to detect the signature of recurrent sweeps may be because *C. grandiflora* has relatively low linkage disequilibrium, increasing power.

However, many positively selected alleles may not follow the trajectory of a classic selective sweep. Soft sweeps — adaptation from an allele previously maintained in the population by mutation-selection-drift balance or the simultaneous fixation of multiple independently derived mutations at the same allele — may still increase the replacement to silent divergence ratio, but are expected to have a smaller effect on linked neutral diversity [48-50]. We expect that soft sweeps will also be common in *C. grandiflora* because of its large *N_e_* [50,51]. In addition, adaptation in genes that contribute to polygenic traits is often expected to occur without fixation of new mutations [52], and this will be missed by both of our tests for positive selection. These considerations suggest that both measures of positive selection are conservative and may miss many instances of positive selection acting in the genome.

Our conclusions about the prevalence of selective sweeps in *C. grandiflora* may seem to conflict with our observation that diversity and Tajima’s D are slightly higher at 4-fold degenerate sites than intergenic sites, since frequent sweeps in coding regions should reduce diversity more strongly in sites near and within genes. There are two likely contributors to this discrepancy. First, recurrent sweeps may in fact reduce average diversity in 4-fold degenerate sites and, by using these sites to set neutral expectations, we are underestimating the strength of purifying selection in intergenic regions. Second, because recombination rates are relatively high, and intergenic regions near coding regions relatively small in *Capsella*, the average impact of linked selection may be similar at 4-fold degenerate sites and intergenic sequences.

### Expression level and selection

Highly expressed genes diverge less than genes with low expression in many species [15-17,19,24,53-55]. This pattern could be due to stronger positive selection in low expression genes or stronger negative selection in high expression genes, or both. Our results suggest that variation in divergence rates between high and low expression genes is largely due to increased negative selection on high expression genes compared to low expression genes. This result is consistent with previous studies that have suggested that new nonsynonymous mutations that cause protein mis-folding or mis-interaction will have stronger deleterious effects in high expression genes than low expression genes and that new mutations that cause mRNA mis-folding are under stronger negative selection in high expression genes than low expression genes [23-25]. In addition, our results agree with a similar study in *Medicago truncatula* that found stronger purifying selection on genes that were expressed than on genes that were not expressed [20].

## Methods

### Sampling and sequencing

Population samples for *C. grandiflora* represented a ‘scattered’ sample of one individual per population for twelve populations from across the geographic range in Greece, plus a thirteenth sample that was the product of a cross of two additional populations (Supplementary Table S1). Plants were grown for several months at the University of Toronto greenhouse, and genomic DNA was extracted from leaf tissue using a modified CTAB protocol. Quality control and single-end genomic sequencing were conducted at the Genome Quebec Innovation Centre at McGill University on the Illumina GAII platform. Each sample was sequenced in 2 to 3 lanes and with a read length of 108 bp.

Leaves from 10 of the 13 individuals were collected and flash frozen for RNA extraction using Qiagen’s RNAeasy plant extraction kit. This RNA was sequenced at the Genome Quebec Innovation Centre, on an Illumina GAII platform with one individual per lane, generating single-end 108 bp long reads. The RNA sequence from these 10 individuals was used for the annotation of the *C. rubella* reference genome, as reported in [30], but the raw sequence data was reanalyzed for this study (see below).

### Genotyping

Genomic reads were aligned to the *Capsella rubella* reference genome [30] using the Stampy aligner 1.0.13 with default settings [56]. Sites around indels were realigned using the Genome Analysis Toolkit (GATK) v1.05777 indel realigner [31]. Genotype and SNP calls were conducted using the GATK UnifiedGenotyper with default parameters [57], after aligning and genotyping the median site quality was 89 and the median individual depth across all sites was 34.

To get a rough assessment of genotyping error rates, we conducted Sanger sequencing from nine coding regions in six of our individuals. From a total of 16,389 bp of Sanger sequence, we found 8 differences between Sanger and Illumina genotypes, giving an estimated error rate of 0.00049. Three of these disagreements were due to three segregating bases at a single site, which we excluded in our GATK genotyping protocol. As we suspect several of these disagreements may be due to Sanger sequencing errors, this provides an upper bound estimate of error rate in coding regions, although higher indel rates and repetitive sequence in noncoding DNA may lead to a higher error rate in those regions.

AFSs were generated from counts of sites in the VCF. Invariant sites were excluded from the AFS if (1) the site quality score was below 90, (2) the fraction of reads containing spanning deletions was not 0 (i.e. the ‘Dels’ value was greater than zero), or (3) any individual’s read depth was less than 20 or greater than 60. Additionally, polymorphic sites were excluded, based on filters 1-3, if (4) the most likely genotype of any individual did not have a phred scaled likelihood score of 0, and if (5) the second most likely genotype had a phred likelihood score less than 40. Additionally, entire regions of the genome were filtered out of the analysis if less than 30% of the sites in a 20kb window passed all other filters. This final filter primarily eliminated pericentromeric regions that were highly repetitive, where we were not confident in genotype calls and observed high heterozygosity.

Our data showed evidence of identity by descent (IBD) in some samples (Supplementary Figure S4). We identified these regions by splitting the genome into 200kb windows, then calculating F_IS_ (Supplementary Figure S5). If F_IS_ was greater than 0.5, the region was flagged as IBD. Across all samples no more than 3 of these regions overlapped. For further analyses we downsampled data in other regions down to 23 chromosomes treating any region of IBD as haploid to ensure that no IBD region was sampled twice from the same individual.

### Divergence

We calculated lineage-specific divergence in two ways. First, we aligned the *C. rubella* reference sequence with sequence data from *Arabidopsis thaliana* and *Neslia paniculata* using lastZ [58] with chaining, as previously described [14]. In order to get an estimate of divergence unique to the *Capsella* lineage, we called sites as diverged where *A. thaliana* and *N. paniculata* had the same nucleotide and this nucleotide differed in the *C. rubella* sequence. If any of the three species was missing data at a site, then that site, and sites 5 bp upstream and downstream of the site, were excluded from divergence analyses in order to avoid inflating divergence because of spurious alignments around indels.

We used a second method for calculating divergence for comparisons that included only coding sequences, particularly for the comparison of genes with different expression levels. We found orthologs between *C. rubella, A. thaliana* and *N. paniculata* genes using InParanoid [59] and MultiParanoid [60]. The peptide sequences of these orthologs were aligned using DialignTX [61], and reverse-translated into coding sequence. Whole-gene divergence at synonymous and nonsynonymous sites was calculated, using PAML [62], under a model where ω was allowed to vary in the *Capsella* lineage compared to other branches.

We conducted comparisons of estimates of the distribution of fitness effects using the two methods above with identical gene sets, and found a very strong concordance of results (data not shown). Furthermore, while we don’t predict a significant effect on results, it is important to note that the two methods also differed in how selected and nonselected classes were determined: the first distinguishes between 0-fold and 4-fold sites and discards other sites, while the second distinguishes between synonymous and nonsynonymous sites, including all data. However, both approaches gave comparable estimates of positive and negative selection (data not shown).

### Identifying conserved noncoding sequences

Conserved noncoding sequences (CNS) were identified in the *C. rubella* genome by first obtaining whole-genome multiple alignments, using a variant of the lastZ/Multiz pipeline previously described [14,63] and using *C. rubella* as the reference genome. The *Capsella* genome sequence was then neutralized (bases replaced with N) and the PhastCons tool used to quantify family-wide levels of conservation. CNSs were then identified, based on extended (>12nt) near-continuous regions of high conservation as previously described [14].

### Estimates of the distribution of fitness effects and α

Site categories were determined based on the Joint Genome Institute’s gene annotation of the *C. rubella* reference genome [30]. The allele frequency spectra (AFS) and divergence values were calculated for each category of sites, and DFE-alpha [28,64] was used to estimate the fraction of sites under negative selection and α, using 4-fold degenerate sites as the neutral reference. The genome was broken up into 10 kb regions and these regions were bootstrapped 200 times to generate 95% CIs for selection on each category of sites. We tested for a significant difference in selection between the pooled set of CNSs and each individual category of CNSs using a randomization test, as in Keightley & Eyre-Walker [28], by calculating the proportion of bootstraps where selection was higher in the pooled set of CNS versus the category of interest. Because this is a two-tailed test, we report twice this proportion as the *p* value.

### Test for signatures of recurrent selective sweeps

We used the multiple species alignments of orthologous genes, generated as described above, to identify silent and replacement single-nucleotide sites that were the same in *A. thaliana* and *N. paniculata* but differed in the *C. rubella* reference, suggesting that the substitution had most likely occurred in the *Capsella* lineage after divergence from *N. paniculata*. From these substitutions, we identified those that did not diverge between *C. rubella* and *C. grandiflora* and were fixed in *C. grandiflora*.

We calculated neutral diversity in sliding windows around fixed substitutions by calculating the proportion of 4-fold degenerate sites within these windows that were polymorphic in *C. grandiflora* (i.e., the proportion of segregating sites). Neutral divergence was measured by calculating the proportion of 4-fold sites within these windows that diverged in the *Capsella* lineage. Diversity/divergence was calculated by dividing diversity by divergence in each window. We conducted this analysis for windows of 500bp, 1kb, and 2kb, extending 40kb from each substitution. We chose this window size range to match analysis done in Sattath et al [39]. For each of the above measures, we bootstrapped by substitution (n=1000) and removed the top and bottom 25 bootstraps to construct 95% confidence intervals. Following Hernandez and colleagues [47], we tested the null hypothesis that diversity/divergence around replacement and silent substitutions does not differ by calculating a one-tailed p value for each window, equal to (*i*+1)/(*n*+2) where *i* is the number of bootstraps in which diversity/divergence around silent sites is lower or equal to the actual diversity/divergence around replacement sites, and *n* is the total number of bootstraps.

To detect linked selection in noncoding DNA, we compared diversity around fixed substitutions within CNSs to diversity around fixed substitutions in non-conserved intergenic regions. To find these substitutions, we compared the multiple sequence alignments of the CNSs between *C. grandiflora, N. paniculata*, and *A. thaliana* and chose sites that differed between *C. grandiflora* and the other species and were fixed within *C. grandiflora*. Additionally, we compared neutral diversity around CNSs with at least one fixed substitution to neutral diversity around CNSs without any fixed substitutions.

### Gene expression

Illumina sequencing generated 331,629,531 reads for 10 individuals, ranging from 31,267,774 to 35,552,133 reads per individual. This RNA sequence was mapped to the *C. rubella* reference genome using Tophat 1.2.0 [65], and expression level was quantified from these mapped reads using Cufflinks 1.3.0 [66]. Cufflinks standardizes expression levels by gene length and library size, returning values in units of ‘fragments per kilobase of exon per million fragments mapped’ (FPKM). We calculated the mean expression level for each gene across our 10 samples and removed those genes with <1 FKPM to eliminate genes that may have been mis-annotated. The remaining 11,564 genes were divided into four, roughly equally sized categories based on expression level: low (1-6.8 FPKM), mid-low (6.8 – 17.5 FPKM), mid-high (17.5-44.7 FPKM), and high (44.7 – 17,092 FPKM). The distribution of fitness effects, α, and ω were calculated for each gene set, using the same protocol described above. We bootstrapped each gene set by sampling genes with replacement 1000 times to generate 95% confidence intervals for selection strength. Using the same methods described for tests of differences within the CNSs categories above, we tested for a significant difference in selection strength between high and low expression genes.

## Acknowledgements

We thank Peter Keightley and Dan Halligan for advice and custom scripts, Yunchen Gong and Emilio Vello for technical assistance, Tanja Slotte, Kate St. Onge, and John Paul Foxe for collecting seeds, and Detlef Weigel, Dan Koenig, Thomas Bureau, Alan Moses, Daniel Schoen, and John Stinchcombe for helpful discussion and/or comments on the manuscript. We would also like to thank Jeff Ross-Ibarra and two anonymous reviewers for helpful comments on the manuscript.

---

Figure S1

Coverage after filtering, across the genome.

A) The number of annotated sites in each category across the genome (light grey), and the number of sites that pass our filters and were used in analysis (dark grey). B) Proportion of sites that pass filters, calculated in 200kb windows, as a function of genomic position.

Figure S2

Pairwise diversity and divergence at 4-fold degenerate sites across the entire genome.

The x-axis represents position along the genome. Statistics were calculated in windows of 5,000 SNPs. Individual lines alternating between grey and blue represent chromosomes. The location of the centromere on each chromosome is indicated by the grey box along the x-axis.

Figure S3

Coding density versus 4-fold degenerate diversity across the genome.

Each point represents one 10 kb window. Black points represent windows that do not overlap centromeres while grey points represent windows that do overlap centromeres. There is a slight negative correlation between diversity and coding density both with and without centromeric windows

Figure S4

Regions of identity by descent in each sample.

The ratio of heterozygous to homozygous calls at sites that are polymorphic across individuals (in 200kb windows) plotted against position across the genome. Each sample is plotted separately. Individual lines alternating between grey and blue represent chromosomes. Regions of IBD were defined as windows where F_IS_ was greater than 0.5 and are indicated by black lines along the x-axis. At most 3 regions of IBD overlap across all individuals. This occurs near the end of chromosome 1.

Figure S5

F_IS_ in windows across the genome in each sample.

F_IS_ in 200kb windows is plotted across the genome. Each sample is plotted separately. Individual lines alternating between grey and blue represent chromosomes. Regions of IBD were defined as windows where F_IS_ was greater than 0.5 and are indicated by black lines along the 0 line of the y-axis.

Figure S6

DFE-alpha results using all alleles, including IBD regions.

The distribution of fitness effects for 0 fold degenerate, 3’ and 5’ UTR, intronic, and intergenic sites are shown. For this analysis the genotyping calls were filtered as described in the methods, but the data was not downsampled in regions of IBD identified in Supplementary Figure S4.

Figure S7

Estimates of positive and negative selection in different categories of CNSs.

A) Distribution of fitness effects. Stars indicate categories in which the fraction of nearly neutral sites was significantly different from the pooled sets of CNSs by a randomization test. B) α and C) ω for each category. Error bars indicate 95% CIs from 200 bootstraps.

Figure S8

Robustness of sweep analysis to different window sizes.

This panel shows the results of our scans for recurrent selective sweeps using alternative window sizes: 500 bp on left and 2kb on right. Otherwise, the methods are the same as described previously.

Figure S9

Additional diversity and divergence data for sweeps around substitutions in conserved noncoding regions.

The left panels show 4-fold degenerate diversity and 4-fold degenerate divergence around substitutions in conserved non-coding sequence (black lines) and non-conserved intergenic sequence (gray shading represents 95% confidence intervals). The right panels show the same information for 4-fold degenerate diversity and divergence around conserved noncoding sequences containing fixed substitutions (black lines) and conserved noncoding sequences without fixed substitutions (gray shading represents 95% confidence intervals).

Figure S10

Allele frequency spectra of replacement sites in genes with different expression levels.

Table S1

Sampling locations of each sample.

Table S2

Allele frequency spectra, summary of diversity statistics, and DFE-alpha model parameters for each site category.

